# The retinoic acid receptor regulates development of a key evolutionary novelty - the molluscan shell

**DOI:** 10.1101/2025.03.09.642214

**Authors:** Keisuke Shimizu, Paul A. O’Neill, Kazuyoshi Endo, Tetsuhiro Kudoh

**Affiliations:** JAMSTEC 2-15, Natsushima-cho, Yokosuka-city, Kanagawa, 237-0061, Japan; Biosciences, University of Exeter, Stocker Road, Exeter EX4 4QD, UK; Department of Earth and Planetary Science, The University of Tokyo, 7-3-1 Hongo, Tokyo 113-0033, Japan

## Abstract

The shells of molluscs are iconic structures of invertebrate exoskeletons supporting and protecting their soft body parts. The shell matrix molecules are synthesized and secreted from a shell-producing tissue, the shell gland. Shell gland cells develop at the early trochophore stage of the larvae. To date, the molecular signalling pathways by which the shell gland forms and starts to secrete the shell remains elusive. Here we demonstrate in the Pacific oyster *Crassostrea gigas* and the limpet *Nipponacmea fuscoviridis*, that the retinoic acid receptor (RAR), is crucial for inducing shell gland formation. RAR is expressed in both species in the shell gland at the late gastrula to early trochophore stages prior to the first production of shell. Suppression of the RAR by chemical inhibitors or gene-knock-down lead to a complete loss of the larval shell. Transcriptomic and *in situ* hybridisation analyses revealed that the developmental regulatory genes that are normally expressed in the shell gland, including *engrailed*, are down-regulated in the RAR-suppressed embryos. Using the RAR functional assay carried out on zebrafish embryos, we also revealed that the oyster RAR cannot transduce the RA signal in zebrafish, indicating that the molluscan RAR is clearly different from vertebrate RARs in its binding capacity to the RA. Our finding represents a key example of adaptive evolution of developmental “toolkit” genes for the origin of a major novel trait, the molluscan shell, in animals.

## Introduction

Fossil evidence, combined with phylogenetic relationships, suggests that skeletogenesis evolved many times in the early Cambrian, accompanying the evolution of novel animal body plans. At this time, molluscs evolved to have one or more mineralized shells. During the process of spiral cleavage in their early development, cells fated to be the shell gland are segregated (Lambert 2010; Perry 2015): by invagination at the gastrula stage, and then the gland is evaginated to form the shell field (Reviewed in Kniprath 1981). Some transcription factors are known to be expressed in the shell gland and shell field cells. Most notable is the homeobox gene *engrailed*, which is expressed in those cells in a specific manner in a variety of species representing major molluscan lineages (Jacobs et al., 2000; Wanninger & Haszprunar 2001; Nederbragt et al., 2002; Hinman et al., 2003; Samadi et al., 2009; 2012; Salamanca-Díaz et al., 2021), suggesting that it may act as a master gene, controlling initial larval shell development. But direct evidence that *engrailed* is responsible for shell formation is lacking, and the genes upstream of *engrailed* in this regulatory network are still unknown. It has also been reported that the Dpp signalling pathway has a crucial role in shell development by regulating shell growth from the early larval stage (Iijima et al., 2008; Kin et al., 2009; Shimizu et al., 2011, 2013; Hashimoto et al., 2012,). In *Lymnaea stagnalis*, asymmetrical expression of Dpp in the left and right side of the shell gland determines asymmetric shell growth, leading to the formation of either dextral or sinistral coiling of the shell, whereas equally distributed Dpp in the left and right sides of shell gland in the limpet *Nipponacmea fuscoviridis* results in L-R symmetric shell growth (Shimizu et al., 2013). Recently, it has been shown that the Wnt signalling pathway is involved in the coiling pattern and morphogenesis of *L. stagnalis* shell (Ohta et al., 2024). Although these studies improved our understanding of shell growth mechanisms, it is still unclear what signals and mechanisms trigger initial shell formation, namely the specification of the lineage of the shell gland and shell field cells.

Transcription factors and nuclear receptors (NRs), bind directly to DNA and regulate the expression of key genes involved in early development, homeostasis, and metabolism. NRs consist of two domains, DNA binding domain (DBD) and ligand binding domain (LBD). LBD in a NR interacts with specific ligands that can activate the NR. Subsequently, the DBD of NRs binds to a specific transcriptional regulatory element that exists in the promoter region of the target genes and regulate their transcription (Germain et al., 2006). The retinoic acid receptor (RAR) is one of the NRs, and its LBD binds to the ligand retinoic acid (RA) in vertebrates. RAR signalling is one of the major signalling pathways that regulates the expression of key developmental genes, including hox genes, during early embryonic development and determines body patterning and tissue differentiation (Marshall et al., 1994, Kudoh et al. 2002). RA is a metabolite of vitamin A (retinol) and mainly synthesized from retinal aldehyde by aldehyde dehydrogenase 1a (Aldh1a) (Kudoh et al. 2002). RA is degraded to oxidized retinoic acid (inactive form) by the RA-degrading enzyme cytochrome P450 26 (Cyp26) to remove unwanted RA molecules (e.g. Kudoh et al. 2002). Hence, these two enzymes, Aldh1a and Cyp26, control the spatio-temporal distribution of RA levels during embryogenesis in chordates (e.g. Niederreither et al., 2002; Kudoh, 2002; Reijntjes et al., 2005).

Here we report that *rar* is predominantly expressed in the shell gland of trochophore larvae of the Pacific oyster *Crassostrea gigas* and the limpet *Nipponacmaea fuscoviridis*, and the loss of function of RAR causes loss of shell gland and shell field, leading to the failure of larval shell formation. Based on these findings, we discuss the master role of the *rar* gene in molluscan shell development and its implications on the evolutionary origin of molluscan shells.

## Results

### Expression of the retinoic acid receptor gene in the shell field cells

To explore the roles of RAR in molluscs, spatial expression patterns of the RAR gene (Cg-*rar*) in the Pacific oyster *C. gigas* larvae were examined using wholemount *in situ* hybridisation. Cg-*rar* is expressed in the dorsal invaginated region, namely the shell field invagination (SFI) (Eyster and Morse, 1984) at late gastrula stage (8 hours post-fertilization [hpf]) (Fig. 1a). The expression of Cg-*rar* in SFI is confirmed by comparison with the expression of two shell field marker genes, Cg-*gata2/3* and *larval shell matrix protein 1* (Cg-*lsmp1*) in SFI. At late gastrula stage (8 pfh), Cg-*gata2/3* is expressed around the SFI (Fig. 1b), but the larval shell matrix protein Cg-*lsmp1* is not expressed yet (Fig. 1c). The expression of Cg-*lsmp1* was first detected in the shell field cells that originated from SFI at early trochophore stage (10 hpf) and overlapped with the Cg-*rar* expression (Fig. 1d, e). Cg*gata2/3* continued its expression around SFI in 10 hpf larvae (Fig. 1f). After the shell field evagination (12 hpf), Cg-*rar* and Cg-*lsmp1* are expressed in the shell field cells (Fig. 1g, h), and Cg-*gata2/3* expression is restricted to the edge of the shell field cells (Fig. 1i). We also examined expression of *rar* (Nf-*rar*) and two shell marker genes, *chitin synthase 1* (Nf-*CS1*) and *6D12,* that were identified as biomineralization genes in the abalone *Haliotis asinina* (Jackson et al., 2006) in the trochophore larvae of the limpet *N. fuscoviridis.* Nf-*rar* is expressed in the shell field cells (Fig. 1j), which are marked by the shell marker gene *6D12* (Fig. 1k). The other shell marker Nf-*CS1* is expressed at the edge of the shell field cells (Fig. l). These observations suggest that *rar* has a role in molluscan larval shell gland development.

**Figure 1.**
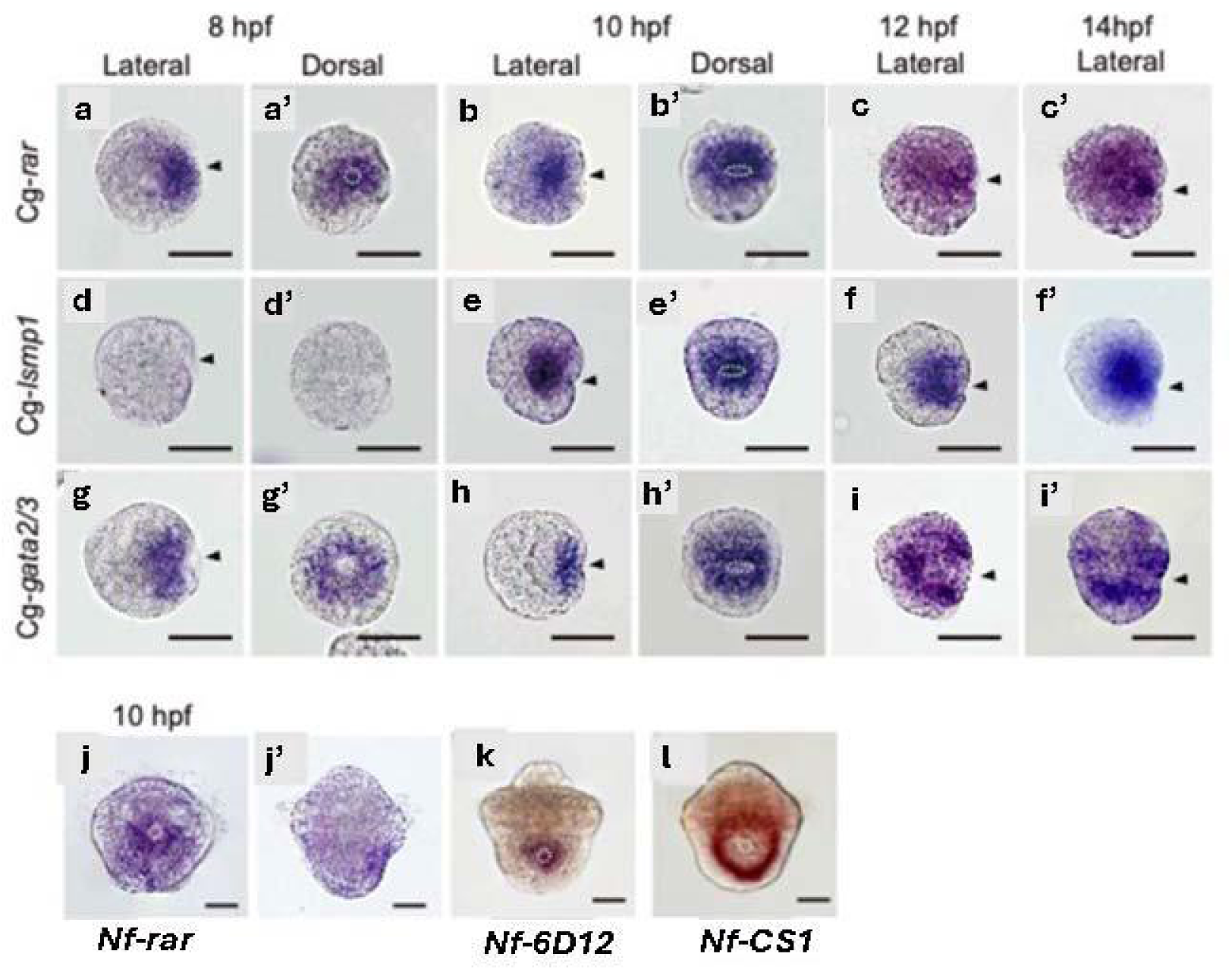
RAR is expressed in the shell gland and progenitor cells from gastrula to trochophore stages. *In situ* hybridisation of the oyster *C. gigas* (a to i) and the limpet *N. fuscoviridis* (j to l). The embryos and larvae were stained with *in situ* probes for *Cg-rar* (a-c), *Cg-lsmp1* (d-f), *Cg-gata2/3* (g-i), *Nf-rar* (j), *Nf-6D12* (k) and *Nf-CS1* (l).

### Retinoic acid receptor is essential for shell fate specification in molluscs

To investigate the function of RAR in molluscs, the embryos of the pacific oyster *C. gigas* and the limpet *N. fuscoviridis* were treated with atRA and two RAR inhibitors (Ro-41-5253 and AGN-193109) at the 1-2 cell stage onward. Although the 24 hpf larvae precipitated the embryonic shell in normal development, the larvae that were treated with the RAR inhibitors completely failed to form their shells both in *C. gigas* (Fig. 2c,d,k) and *N. fuscoviridis* (Fig. 2h,i,l). The 24 hpf larvae treated with at-RA showed abnormal morphology of larval shells with reduction of areas covered by the shell (Fig. 2b,g). We then confirmed the crucial role of RAR in the shell development using another loss-of-function approach: Nf*rar* was knocked down by microinjection of a specific morpholino oligo (Nf*rar*-MO) in *N. fuscoviridis*. The Nf*rar*-MO injected embryos indeed developed non-shelled veliger larvae (Fig. 2n), which are similar to those treated with RAR inhibitors (Fig. 2X). These results suggest that RAR has a crucial role in promoting shell development in molluscan larvae.

**Figure 2.**
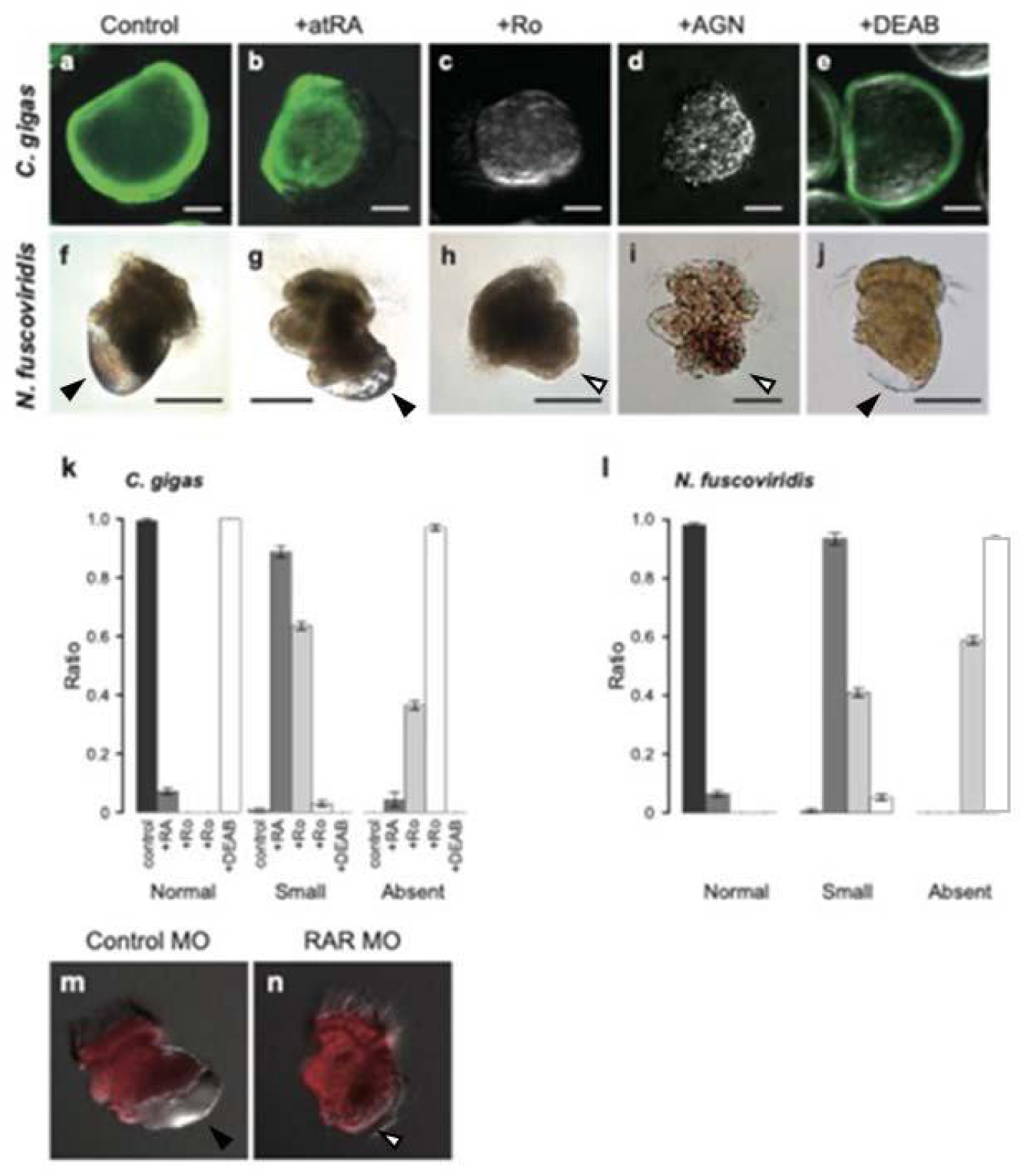
Suppression of RAR blocks shell formation in the oyster and limpet larvae. Live *C. gigas* (a-e) and *N. fuscoviridis* (f-j) embryos were treated with atRA (b,g), the retinoic acid inhibitors RO (c,h), AGN (d,i) and aldh1a inhibitor DEAB (e,j). For *C. gigas*, the live larval shell was visualized by calcein with green fluorescence (a-e) indicating absence of the shell in the Ro and AGN treated embryos (c,d). In the limpet, presence (black arrow head) and absence (white arrow head) of the larval shell was visible with bright field microscope (f-j). k,l. Ratios of normal shelled larvae, small shelled larvae, and abnormal larvae in three biological replicate experiments using the oyster (**k**) and the limpet (**l**). Scale bar: 20μm (**a**) and 50 μm (**b**). (m.n.) Gene knockdown of *Nf_rar* by morpholino injection. Consistent with the results of Ro and AGN, Nf*rar*-MO suppressed larval shell formation in the limpet (n, white arrow head).

### RNAseq reveals that RAR regulates the shell field genes

In order to identify the target genes for the retinoic acid receptor (RAR) in molluscs, we treated the oyster, *C. gigas* embryos/larvae with RAR inhibitor (Ro-41-5253) and atRA at the 1-2 cell stage onward and analzyed the transcriptome in early trochophores (9.5 hpf). The results show that the expression levels of 1289 and 426 genes (14.1 and 4.7 %) were significantly changed by the Ro-41-5253 and atRA treatment, respectively (|logFC| > 1 and FDR < 0.05, Fig. 3a). If Ro-41-5253 blocks the atRA signal, a common set of genes would be expected to show an inverted response to these treatments. However, we found only one gene that was down- and up-regulated by Ro-41-5253 and atRA respectively. This suggests that atRA may not act as the ligand of RAR to activate the genes downstream of RAR. In contrast, 246 and 47 genes were down- and up-regulated in both Ro-41-5253 and atRA treated larvae, respectively (Fig. 3a), suggesting there may be some level of similarity between atRA and Ro-41-5253 treated embryos. Indeed, a LogFC plot of these two conditions shows a linear correlation pattern through the down-regulated to up-regulated genes (Fig. 3b). It should also be noted that the angle of the plot leans toward the Ro-41-5253 side, suggesting the level of gene expression change is more prominent in the Ro-41-5253 compared with atRA treated samples (Fig. 3b).

**Figure 3.**
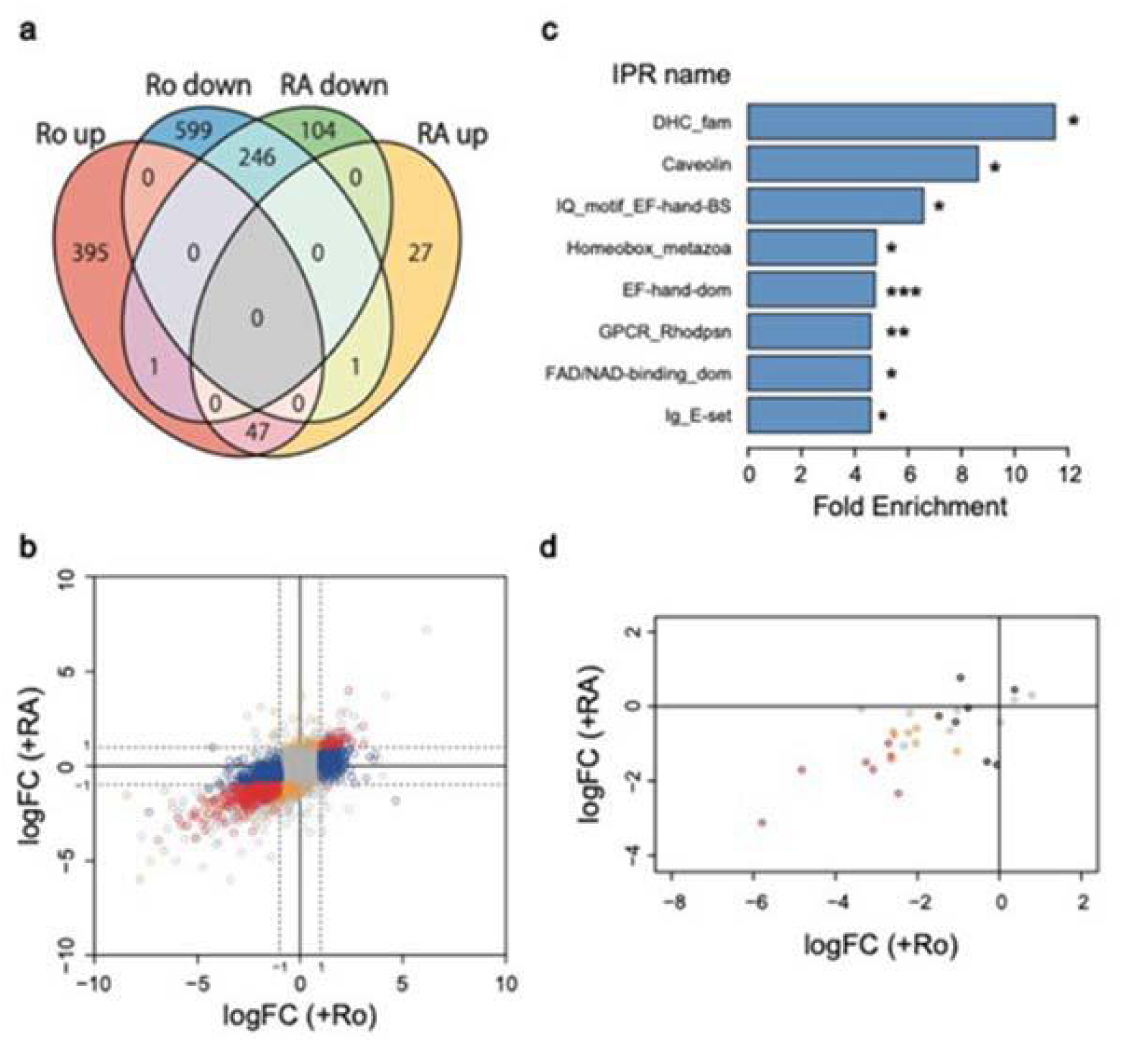
Transcriptomics analyses of *C. gigas* 24hpf larvae treated with RA or Ro. Number of transcripts up or down regulated by RA and/or Ro (|logFC| > 1 and FDR < 0.05. LogFC plot of expression levels of RA and Ro samples at Y and X axes, respectively. Red highlights up- or down-regulated genes by RA. InterProScan enriched gene families in the transcripts down-regulated by Ro identified 8 gene familes/domains. d. LogFC plot of expression levels of RA and Ro samples focusing on homeobox genes.

Among the 846 genes that were down-regulated in Ro-41-5253 treated larvae (Fig. 4a), eight key domains or families are significantly enriched in InterProScan (dynein heavy chain domain, caveolin, EF-hand domain, homeobox, G protein-coupled receptor, FAD/NAD (P) binding domain, and immunoglobulin) (Fig. 4c) (p < 0.05). We then focused on the larval shell matrix protein (LSMPs) encoding genes in *C. gigas* (Zhao et al., 2018). We found that a total of 13 and 9 of the 29 Cg-LSMPs encoding genes were significantly down regulated in Ro-41-5253 and atRA treatment, respectively (Fig. 3b). These genes had relatively high expression levels in the control samples (FPKM > 10) except for one gene, CGI_10008969, in Ro-41-5253 (Fig. 3b). The data also confirmed that the effect of Ro-41-5253 is stronger than that of atRA. The homeobox genes *engrailed*, *post2, evx*, *cux*, and *ssuh2,* were down-regulated in both conditions (Fig. 3d), and other homeobox genes, *hox1, lox1, hhex*, *nkx6.2*, *pou4*, *six1/2* and *rx,* were down-regulated only in Ro-41-5253 treatment, suggesting these genes might be involved in shell gland and shell field cell differentiation and function (Fig. 3d).

**Figure 4.**
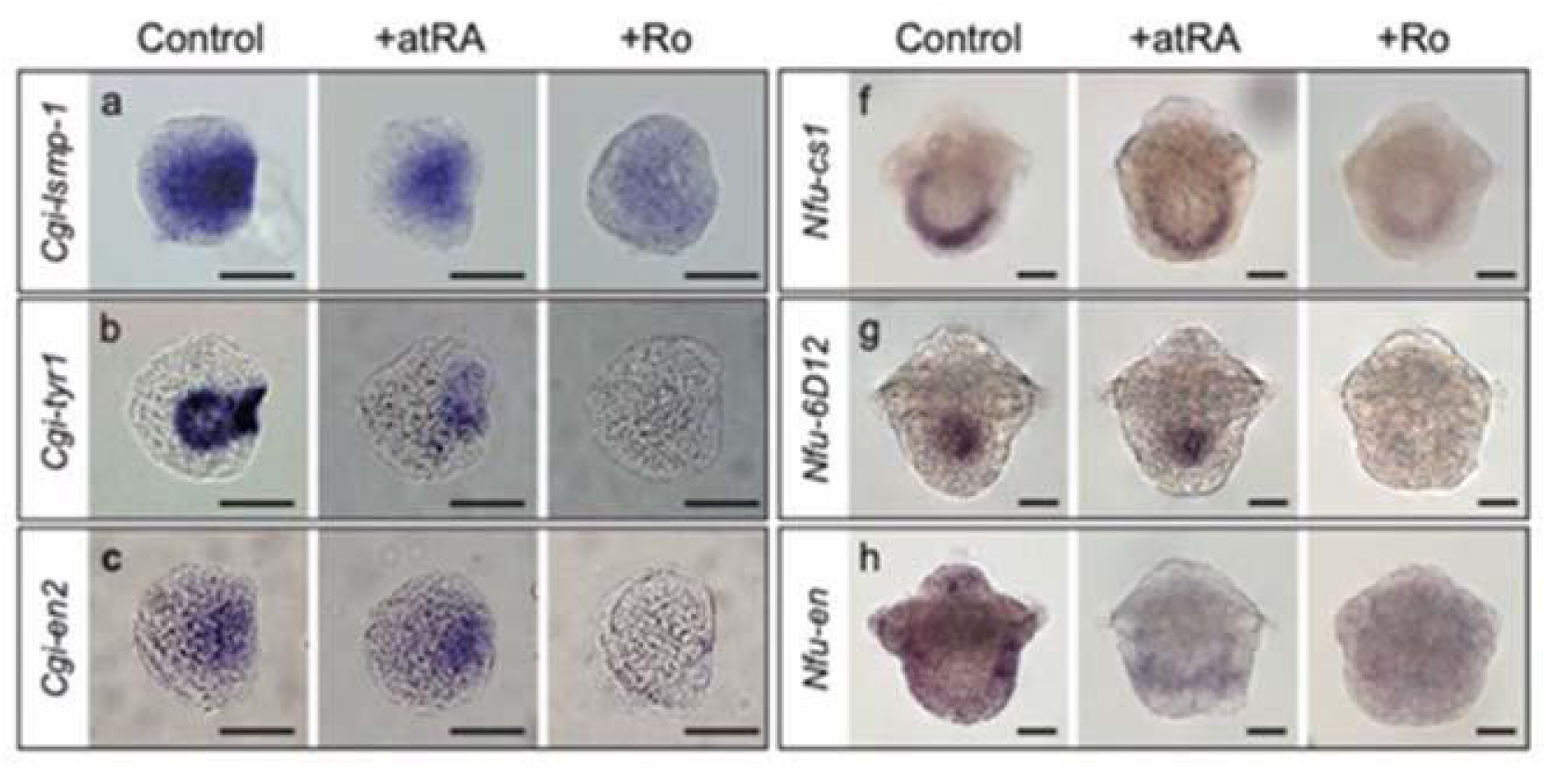
Gene expression patterns in shell gland and shell field are dependent on RAR signalling. *In situ* hybridization of oyster (18hpf) (a-c) and limpet (f-h) (10h) larvae. Expression of *Cg-lsmp-1* (a), *Cg-tyr1* (b), *Cg-en2* (c), *Nf-cs1* (f), *Nf-6D12* (g) and *Nf-en* (h) are all suppressed by the RAR inhibitor (Ro-41-5253). Embryos treated with all trans Retionic Acid (atRA) did not show strong up or down regulation of these marker genes. Scale bar: 20μm.

### RAR determines shell gland cell fate

To confirm that shell-related gene expression is regulated by the RAR signalling pathway, Ro-41-5253 treated larvae were *in situ* stained with probes for shell-marker genes (Pacific oyster: Cg*engrailed,* Cg-*lsmp1* and Cg-*tyr1*; limpet: Nf-*engrailed,* Nf-*CS1* and Nf-*6D12*) in *C. gigas* (8 hpf trochophore larvae) and *N. fuscoviridis* (10 hpf trochophore larvae) (Fig. 3a, b, e, f). Ro-41-5253-treated larvae showed suppressed expression of these shell-marker genes in both species, strongly suggesting that RAR signalling has a crucial role in cell fate specification of the shell gland. Besides these shell field-marker genes, the transcription factor *engrailed* was also suppressed in the shell gland region of both species after Ro-41-5253 treatment.

### CgRAR disturbs the RA signaling in zebrafish embryo

The DNA binding domain (DBD) of oyster RAR (CgRAR_DBD) has a high similarity (81-86%) to that of chordate RAR. In contrast, the ligand binding domain (LBD) of oyster RAR (CgRAR_LBD) shows a low similarity (53-60%) to that of chordate RAR. It has been known that the 25 amino acid sequence around the LBD region, called the ligand binding pocket (LBP), is structurally important for interaction with RA in the human RAR. Five of these 25 amino acids in the LBP of CgRAR are different from human and zebrafish RAR suggesting that the ligand-binding ability and/or specificity might be different between CgRAR and vertebrate RARs. To test if Cg-RAR could respond to the RA *in vivo*, we used zebrafish (*Danio rerio*) embryos as an *in vivo* assay system. Firstly, zebrafish RARab (Dr*RARab*) was overexpressed in the zebrafish embryo by injecting mRNA at 1-cell stage. When uninjected zebrafish embryos were treated with 0.1 µM atRA, they showed mild reduction of the head and tail (Fig. 5Bb). However, when the Dr*RARab* was overexpressed, the head and tail became further reduced (Fig. 5Be), suggesting that overexpression of Dr*RARab* enhanced the action of atRA and induced the posteriorisation of the brain (Kudoh et al. 2002). However, when Cg*RAR* is overexpressed and the embryos subsequently treated with 0.1 µM atRA, the phenotype was similar to that of uninjected embryos treated with 0.1 µM atRA (Fig. 5Bk), suggesting that CgRAR did not properly transduce the signal of atRA in the zebrafish embryo. To examine if the lack of response is due to the different LBD of the CgRAR, we generated a chimeric construct of DrRARab in which the LBD was replaced with that of CgRAR (DrRAR-CgLBD) (Fig. 5A). Indeed, the embryos injected with mRNA of DrRAR-CgLBD did not show any enhancement of the effect of 0.1 µM atRA (Fig.5Bh). The same result was observed with 9-cis-RA, indicating that CgRAR cannot respond to either at-RA or 9-cis-RA as the DrRAR does (Fig. 5Bc,f,i). To further examine brain-regional specific action of the three RAR constructs (DrRAR, CgRAR, and DrRAR-CgLBD) to atRA, we conducted expression analysis using three positional marker genes, Dr*pax2* (mid-hindbrain boundary [MHB]), Dr*hoxb3a* (tail to hindbrain rhombomere 5) and Dr*hoxb5b* (tail to hindbrain rhombomere 7). Uninjected embryos treated with 0.1 µM atRA showed loss of Dr*pax2* and expansion of Dr*hoxb3a* and Dr*hoxb5b* to the anterior end (Fig. 5Cb,f,j), suggesting the loss of the forebrain, midbrain and MHB. However injection of the Cg-*rar* mRNA rescued the Dr-*pax2* expression in the MHB and moved the anterior border of the Dr-*boxb3a* and Dr-*hoxb5b* expression domain backward in the presence of 0.1 µM at-RA (Fig.5Cd.h.j), suggesting Cg-RAR acts as a dominant negative form against the Dr-RAR in the zebrafish embryo. Consistently, the DrRAR-CgLBD showed a similar rescue phenotype (Fig.5Cc,g,k), indicating the dominant negative activity is due to the diverse functions of the LBDs in the oyster and zebrafish RARs. We then examined Dr-*hoxb1b* expression, one of the direct target genes of RA signaling, using zebrafish embryos at late gastrula. DrRAR-CgLBD or Cg*RAR* mRNA injection led to suppress Dr-*hoxb1b* expression (Fig. 5Db,c).

**Figure 5.**
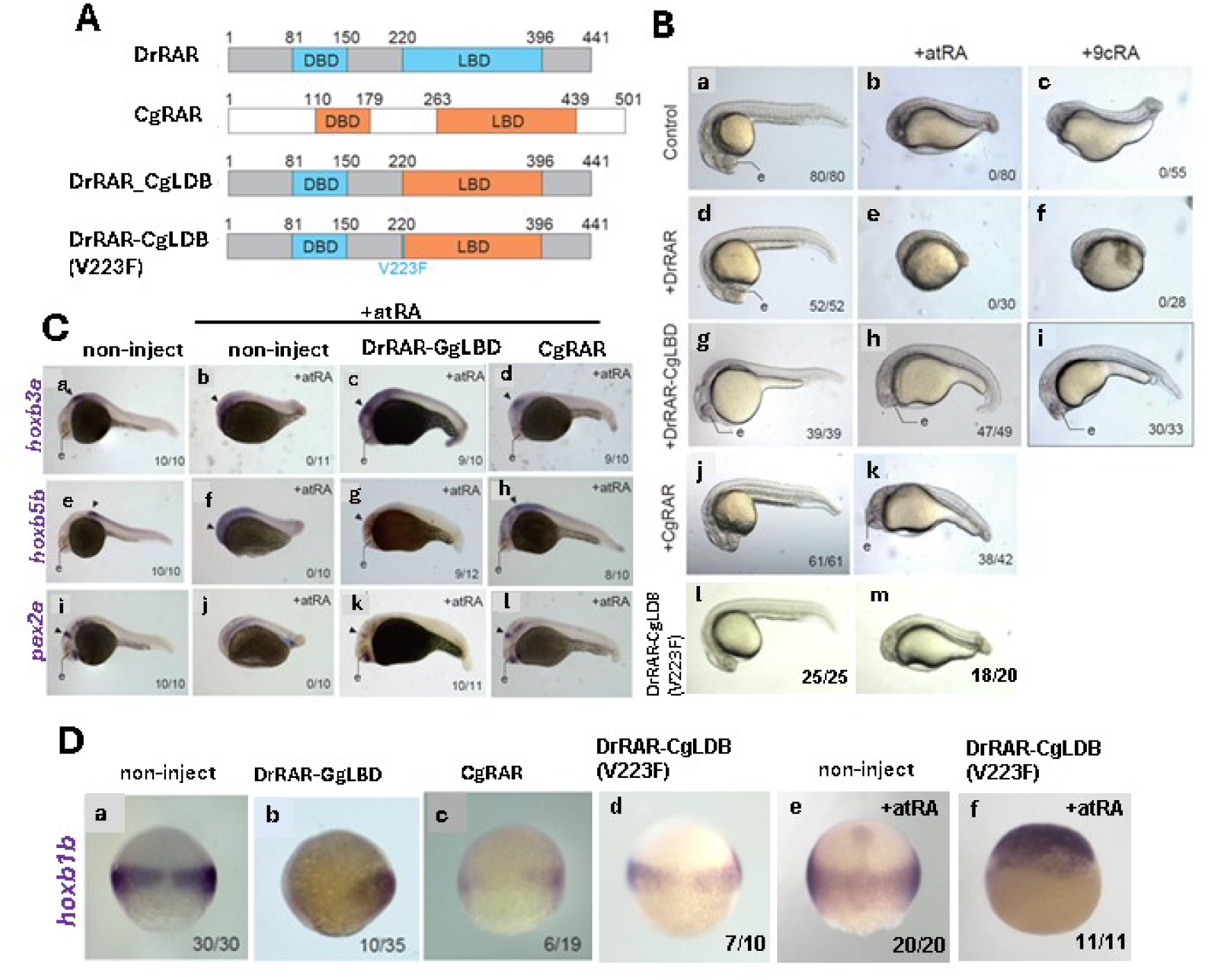
Comparative and functional domain analysis of *Cg-rar* and *Dr-rar* using the zebrafish embryo *in vivo* model. A. zebrafish RAR (DrRAR) and Pacific oyster RAR (CgRAR) show conserved domain structure with N-terminal DNA binding domain (DBD) and C-terminal ligand binding domain (LBD). Chimeric construct of DrRAR_CgLDB (DrRAR backbone replaced with the LBD domain of CgLBD), and DrRAR_CgLDB(V223F), a point mutation of DrRAR_CgLDB that altered the amino acid V(valine) 223 to F(phenylalanine), B. Fertilised zebrafish eggs were injected with mRNA encoding RAR constructs listed in A and treated with or without atRA or 9cRA. Live images were taken at 24hpf. C. *In situ* hybridisation of 24hpf embryos injected with RAR constructs, treated with atRA, and stained with *Dr_hoxb3a* (a-d), *Dr_hoxb5b* (e-h) or *Dr_pax2a* (i-l) probes. D. *In situ* hybridisation staining of *Dr-hoxb1b* in the injected and RA treated embryos. Embryos in b and c were injected with mRNAs at 2-cell stage in one blastomere, showing effects in one side of the embryo only.

Finally, to identify the key amino acid that alters LBD activity between vertebrates and molluscs, the amino acid Valine (V) at position 223, a residue that is only specific to molluscs, was changed to Phenylalanin (F), which is conserved in vertebrates (DrRAR-CgLBD V223F), its activity was examined. Although DrRAR-CgLBD showed a dominant negative activity, this single amino acid modification DrRAR-CgLBD V223F was sufficient for the gene to lose the dominant negative activity, and act instead similar to the zebrafish RAR, enhancing the RA treatment in the 24hpf embryos by reducing head and tail (Fig. 5Bm) and by inducing *hoxb1b* expression at late gastrula stage (Fig. 5Df). Altogether, we conclude that the molluscan RAR has a property distinct from the vertebrate RAR in response to the RA, and this difference is largely due to the single amino acid substitution at the position 223 with valine in molluscs and phenylalanine in vertebrates respectively.

## Discussion

We report here that the retinoic acid receptor (RAR) is essential trigger for shell gland development and initial shell secretion in molluscs. The expression patterns of the homeobox-containing regulatory gene *engrailed* are likely associated with the development of molluscan shells, as *engrailed* is expressed in the ectoderm cells at the boundary of the shell forming region of various taxa (Polyplacophora, Jacobs et al., 2000; Cephalopoda, Bratte et al., 2007; Scaphopoda, Wanninger and Haszuprunar, 2001; Bivalvia, Kin et al., 2009, Gastropoda, Moshel et al., 1998). Other transcription factors, *distal-less* (*dlx*)*, soxC, gata2/3, pax2/5/8, and lox 4* are also expressed inside or the boundary of the shell field cells in bivalve or gastropod (Jackson and Degnan 2016; Liu et al., 2015, 2017, 2020). However, there has been no functional evidence showing that these transcription factors are involved in the initial shell development. In this study, we found that *engrailed* expression in the shell field was significantly down-regulated in the larvae of both oyster and limpet that were treated by Ro-41-5253 (Fig. 3c, h), and these larvae failed to develop their embryonic shells with down-regulation of the expression of LSMP genes (Figs. 2-4). On the other hand, the expression of other shell-related candidate genes (*dpp*, *dlx*, *gata2/3*, and *soxC*) was not affected under the Ro-41-5253 treatment in oyster larvae (9.5 phf). These results suggest that RAR signaling directly or indirectly regulates *engrailed* expression, and both *rar* and *engrailed* are involved in the differentiation of shell secretary cells and crucial signals for their embryonic shell development. In contrast, other shell related genes (*dpp*, *dlx*, *gata2/3*, and *soxC*) possibly belong to other gene regulatory networks (GRN) and play roles in the shell growth or other shell-related functions rather than the differentiation of shell secretory cells.

The signaling pathway encompassing the retinoic acid receptor has been well studied in chordates. Its ligand, RA, is an important morphogen involved in body axis formation and growth patterning corresponding to its gradient, as RA can diffuse over long distances, and its spatio-temporal distribution regulates the expression of homeotic genes (Marshall et al. 1994; Aulehla & Purquite, 2010). In contrast, RAR is not found in insect species such as *Drosophila,* thus it has been suspected that the morphogenetic role of the RA/RAR signaling may have been acquired along the branch leading to chordates, and therefore is linked to the origin of their innovative body plans (Shimeld 1996; Manzanares et al. 2000; Schilling & Knight 2001; Wada 2001; Holland 2005). However, a recent study using the genomic databases including Ambulacraria and Protostome revealed that the components of RA signaling machinery, RA synthetase (*aldh1a*), RA-degrading enzyme (*cyp26*) and receptors (*rar*), originated in the last common ancestor of bilaterians, rather than in the last common ancestor of chordates (Albalat & Canestro 2009). This suggests that RA signaling pathway may well have been involved in the evolution and developmental diversity of not only Chordata but also Lophotrochozoa. The effects of RA overdose have been studied in a variety of lophotrochozoans including platyhelminthes, annelids, and molluscs (e.g. platyhelminthes: Romero & Bueno, 2001, annelids, Handberg-Thorsager et al., 2018, molluscs: Creton et al., 1993; Dmetrichuk et al., 2006; Carter et al., 2015; Vogeler et al., 2017; Johnson et al., 2019). These previous reports suggest that an ancestral function of RAR signal was neuronal growth and cell survival (Dmetrichuk et al., 2006; Campo-Paysaa et al., 2008; Handberg-Thorsager et al., 2018). In chordates, an important role of RAR signaling is embryonic patterning along the anterior-posterior axis via regulation of *hox* genes (Shimeld, 1996; Schilling and Knight, 2001, Kudoh et al. 2002). For instance, treatment of the zebrafish embryo with atRA induces loss of the anterior brain cell fates (Fig. 5, Kudoh et al. 2002). However, a loss of these anterior structures has not been observed in invertebrate larvae treated with atRA. Our transcriptome results also showed that atRA does not induce expression of *hox* genes (Fig. 3), albeit some hox and other homeobox genes were down-regulated (*hox1*, *lox5*, *lox4*, and *post2*) by Ro-41-5253 (Fig. 3). Interestingly, many homeobox genes that are downregulated by Ro-41-5253 showed an overall linear relationship to the downregulated pattern seen in the treatment with atRA, suggesting that atRA in this experiment may have acted as a weak inhibitor of RAR. If this is the case, it might be possible that molluscan RAR may have a ligand other than atRA (or 9cisRA). Or otherwise, the role of RA in molluscs could primarily be to restrain the activity of RAR. Within the 846 genes that were down regulated in the Ro-41-5253 treated larvae (Fig. 4a), eight key domains or families are significantly enriched in InterProScan (dynein heavy chain domain, caveolin, EF-hand domain, homeobox, G protein-coupled receptor, FAD/NAD (P) binding domain, and immunoglobulin) (Fig. 4c) (p < 0.05). One of the enriched domains, EF-hand domain, is well known in calcium-binding proteins, which may be involved in the secretion of calcium carbonates in the larval shell. Homeobox containing genes play key roles in the gene regulation for fate specification and body patterning. These data suggest that RAR is involved in the process of cell fate specifications including regulation of gene expression for key transcription factors (such as *engrailed* and tissue specific genes including calcium secretion proteins and LSMPs.In chordates, it is known that the RAR/RXR heterodimer binds to the RA response elements (RAREs) and maintains the expression of target genes. RAREs consist of two direct repeats of a core hexameric motif 5’-(A/G)G(G/T)(G/T)(G/C)A-3’, separated by 1, 2, or 5 bp long nucleotides (DR1, DR2, or DR5, respectively) (Balmer and Blomhoff, 2002). In the mollusc *Nucella lapillus* and the annelid *Platynereis dumerilli*, the RAR/RXR heterodimer is able to bind to RAREs (DR1, DR2, and DR5) like chordate counterparts (Gutierrez-Mazariegos et al., 2014; Handberg-Thorsager et al., 2018). Indeed, sequences of DBD in RAR including the three box regions (P-box, D-box, and T-box) are highly conserved between vertebrates and lophotrochozoans. In the zebrafish embryo assay, CgRAR acted like a dominant negative form and antagonized the signaling pathway with DrRAR. This suggests that CgRAR cannot bind to atRA in the manner similar to chordate RAR but can still bind to the RARE DNA sequence via DBD, therefore, can act as a dominant negative form in zebrafish.

Although the DBD of RAR is well conserved among lophotrochozoans and chordates, LBD sequences of RAR in lophotrochozoans are more divergent from those in chordates. In the annelid *P. dumerilli,* a single amino acid difference at position 356 is observed among the important residues of the LBP, at which position human and other vertebrates have phenylalanine while the annelid has valine, and this mutation (V356F) is shown to decrease the binding ability to atRA (Handberg-Thorsager et al., 2018). The valine at position 356 in RAR of *P. dumerilli* is shared by the annelid *Capitella teleta*, pacific oyster *C. gigas*, and the brachiopod *Lingula anatina*. In the zebrafish embryo assay, the phenotype and the gene expression pattern under the overexpression of CgRAR and DrRAR-CgLBD with atRA treatment rescue, indicated a behaviour as an antagonist to DrRAR (Fig. 5). On the other hand, the zebrafish embryos with overexpression of DrRAR-CgLBD-V223F, which corresponds to V356F mutant of *P. dumerili* RAR (Handberg-Thorsager et al., 2018), mimicking vertebrate RAR, indicated results just like those of the overexpression of DrRAR (Fig. 5). Indeed, DrRAR-CgLBD-V223F was able to induce *hoxb1b* expression, a direct target gene of RAR in zebrafish, while DrRAR-CgLBD inhibited its expression (Fig. 5). These results suggest that the RAR of molluscs and other lophotrochozoans such as annelids and brachiopods cannot activate downstream gene expression using atRA as the ligand, and that the single amino acid modification in the LBP critically affects the ligand binding and/or transcriptional regulation. The marginal or differential role of atRA in molluscs is also supported by the experiment with DEAB, an inhibitor of atRA synthesis, showing no clear inhibitory effect in the shell development (Fig. 2).

Invertebrate exoskeletons constitute one of the key products of the Cambrian explosion, and may have diversified by, and contributed to, the evolutionary arms race, because external skeletons play important roles for survival (e.g. feeding, protection, and body support). Major bilaterian lineages with exoskeletons have probably evolved their hard tissues independently in each phylum within a very short time across the Ediacaran-Cambrian transition (Murdock and Donoghue, 2011). However, the evolutionary origins of exoskeletons remain unclear. Various homologous genes or proteins that are associated with skeletogenesis have been reported from different groups of bilaterians (Ettensohn et al. 2003; Livingston et al., 2006; Jackson et al., 2007), leading to a notion that a “biomineralization toolkit” might have evolved in parallel among those taxa. The homeobox-containing transcription factor *engrailed* has been thought to be one of the key biomineralization toolkit genes. Jacobs & Gates (2003) compared expression patterns of *engrailed* among invertebrate phyla and pointed out the relationship between the *engrailed* expression pattern in ectodermal cells and skeletal development among protostomes; absence of *engrailed* expression in ectodermal cells corresponds with absence of exoskeleton in onychophorans (Wedeen et al., 1997) and annelids (Wedeen & Weisblat, 1991; Lans et al., 1993; Seaver & Kanesige, 2006). In juveniles of brittle stars, *engrailed* is expressed in the boundaries between newly forming spine ossicles (Lowe & Wray 1997). The cells that form spine ossicles are called skeletogenic mesenchyme cells, which are derived from mesoderm, but they rely in part on ectodermally derived cues (Malinda & Ettensohn, 1994). Thus, the patterns of *engrailed* expression in skeleton boundaries in echinoderms are consistent with those in protostomes. Similar *engrailed* expression patterns have been reported in Lophotorochozoa. In molluscs and brachiopods, *engrailed* is expressed around the shell gland and the mantle lobes that are derived from ectodermal cells, respectively (Moshel et al., 1998; Jacobs et al., 2000; Wanninger and Haszuprunar, 2001; Bratte et al., 2007; Kin et al., 2009; Shimizu et al., 2017). We found that RAR signal is the key for inducing shell gland development and thus subsequent shell formation, and regulates the expression of the homeobox gene *engrailed*. Thus, previously existing gene modules including *engrailed* that have been deployed in dorsal ectodermal cells might have contributed to molluscan shell evolution, and this novel body plan is most likely to have been recruited by the RAR pathway. Such an evolutionary process, in which a new gene function is added to an existing toolkit, known as co-option, is not limited to the origin of the molluscan shell (Carrol et al., 1994; beetle horn, Moczek & Rose, 2005). Qur new findings provide clues about the origin and evolution of exoskeletons based on the shell gland cell development and involvement of divergent nuclear receptor signalling pathway, and pose more general questions about how various novel body plans evolved in animals.

## Material and Methods

### Animals

Individuals of *Nipponacmaea fuscoviridis* were collected from the intertidal rocky shores in Manazuru, Kanagawa, Japan. Individuals of the oyster *Crassostrea gigas* were obtained from Guernsey Sea Farms Ltd. (Guernsey, UK). Methods of egg collection followed previous studies (oyster: Vogeler et al., 2017, limpet: Deguchi 2007). Embryos of the limpet and oyster were cultured in filtered seawater at 24 °C and 22 °C, respectively.

### Chemical treatment

Embryos of *C. gigas* and *N. fuscoviridis* were incubated in filtered natural seawater containing 0.1% (v/v) DMSO and treated with 2μM of all-trans retinoic acid (R2625, Sigma-Aldrich Japan, Tokyo, Japan) and 0.5-2 μM of RAR inhibitors Ro-41-5253 (SML0573, Sigma-Aldrich Japan, Tokyo, Japan) and AGN-193109 (SML2034, Sigma-Aldrich) at the 1 or 2-cell stages. As a negative control, embryos were also exposed to filtered natural seawater with 0.1% (v/v) DMSO. For the examination of the presence or absence of calcium carbonates in the larval shells of oyster, we added 1/400 calcein stock solution (Sigma, 6g/L in DMSO) into culture dishes from 4 to 24 hpf embryo.

### Microinjection of MO and mRNA

The MO (Gene Tools LLC) NfuRAR-MO (GTCATACATTGTATTTGGCACATCA) was designed against their start codons. NfuRAR-MO was adjusted to 2.0 mg mL^-1^ in 3 % Rhodamine B dextran isothiocyanate-Dextran (R8881, Sigma). Full length of CgiRAR (CGI_10028545) was amplified by PCR using specific primers (CgiLBD-F TGTGGGATAAAGTGACAGAAC, CgiLBD-R CTAACCGTAAATGAAGGACTC, ZfRAR_CgiLBD-F TTCATTTACGGTTAGAGATCCCAGGCTCCATG and ZfRAR-CgiLBD-R CTGTCACTTTATCCCACA-GGTCGACATCCAG) and cloned into pCS2+ vector. The mRNA was synthesized using the mMESSAGE SP6 kit (AM1340, Thermo Fisher Scientific) and adjusted to 25 µg mL^-1^ in 5 % Phenol red (Sigma). NfuRAR-MO was injected into unfertilized eggs of *N. fuscoviridis* using FemtoJet microinjector (Eppendorf). The cocktail of MOs

(DreRARaa and ab) and mRNA (CgiRAR) were injected into one-cell zebrafish embryos.

### RNA extraction and transcriptome analysis

We collected 9 hpf trochophore larvae of oyster that are three conditions in triplicate (0.1% DMSO, 1 µM of all-trans retinoic acid, and 1 µM of RAR inhibitor Ro-41-5253) and 10 hpf trochophore larvae of limpet that are no treatment in triplicate. We isolated mRNA according to the manufacturer’s protocols for RNA extraction using TRIzol reagent (15596026, Thermo Fisher Scientific) and RNeasy (Quiagen, 74104), and stored at −80 °C until for complementary DNA (cDNA) synthesis and/or transcriptome analysis. We prepared 100-bp DNA libraries from the mRNA samples using ScriptSeq RNA-seq library preparation kit (ScriptSeq: V224-40412) according to the manufacturer’s protocols and then performed 100-base paired-end sequencing by HiSeq2500 system (Illumina). We then performed de novo transcriptome assembly using Trinity (Grabherr et al., 2011; Haas et al., 2013) release (r20131110) and mapping these reads using Tophat (version 2.0.8b). We picked up transcripts of which FPKM values are higher than 5 for our analyses. Differential expression analysis was evaluated using edgeR in the bioconductor package of R (3.7.14). To examine the changes of expression levels of larval shell matrix protein (LSMP) genes by Ro and RA, we selected candidate genes from our *C. gigas* larval transcriptome database (Zhao et al., 2018). Among 110 annotated Cgi-LSMPs genes in D-shape larvae of *C. gigas*, we selected 29 Cgi-LSMPs for analyses; we removed 58 housekeeping-like genes that are constantly expressed in all developmental stages (38 stages) and adult tissues (11 tissues) (RPKM > 1, Zhang et al., 2012), 13 transcripts that are not predicted as gene models in the genome study (Zhang et al., 2012), and 11 transcripts that are quite low expression levels in our transcriptome result (FPKM < 1). cDNA synthesis and gene subcloning cDNA was synthesized according to the manufacturer’s protocols for cDNA synthesis using ReverTra Ace (TRT-101,Toyobo, Osaka, Japan). Nine gene sequences (*Cgi-rar*, *Cgi-cyp26, Cgi-engrailed*, *Cgi-gata2/3*, *Cgi-tyr1*, *Cgi-lsmp1, Nfu-rar*, *Nfu-cyp26*, *Nfu-6D12*) were amplified with PCR using specific primers and then purified using the SV Gel Extraction and PCR Clean-Up system (A9281, Promega). Subcloning of these genes was performed using pGEM-T easy Vector Systems (A1360, Promega) and *Escherichia coli* JM109 competent cells (L1001, Promega).

### Whole mount in situ hybridization

Molluscan embryos were fixed with 4% paraformaldehyde (PFA) buffer (with 0.1M MOPS, 0.5M NaCl, and 2mM EGTA) for 1 h at room temperature. After fixation, samples were washed with phosphate-buffered saline with 0.1 % Tewwn-20 (PBT) for 5 minutes five times, followed by the dehydration with a gradual series of methanol/PBT (50/50, 80/20, 100/0) for 15 minutes each, and stored in 100 % methanol at −20°C. Zebrafish embryos were fixed with 4% PFA in PBS at 4 °C overnight, washed three times with PBT, and stored in 100 % methanol at −20°C. Probe synthesis and whole mount *in situ* hybridization was performed following the protocol from a previous study on the brachiopod *Lingula anatina* (Shimizu et al., 2017) and zebrafish *D. renio* (Kudoh et al. 2002) for molluscan embryos and zebrafish embryos respectively. Color development was performed with NBT/BCIP (Sigma-Aldrich, 11681451001) or BM Purple (Sigma-Aldrich, 11442074001) in the dark. All color reactions were stopped at the same time.

**Table 1.** Number of gene expressed-larvae in the oyster and limpet.

***C. gigas* (18 hpf)**
|  | Trial | *tyr* | *smp-L1* | *collagen* | *en* | *cyp26* |
| --- | --- | --- | --- | --- | --- | --- |
| **Control** | 1 | 36/38 | 35/40 | 39/42 | 47/51 |  |
|  | 2 | 50/55 | 32/35 | 40/45 | 39/40 |  |
| **RA (2 μM)** | 1 | 34/40 | 35/39 | 30/37 | 32/40 |  |
|  | 2 | 48/55 | 42/46 | 28/38 | 40/48 |  |
| **Ro (2 μM)** | 1 | 3/35 | 4/50 | 2/40 | 8/44 |  |
|  | 2 | 5/52 | 41/48 | 5/44 | 3/35 |  |

***N. fuscoviridis* (10 hpf)**
|  | Trial | *cs-1* | *6D-12* | *en* | *dpp* | *cyp26* |
| --- | --- | --- | --- | --- | --- | --- |
| **Control** | 1 | 32/32 | 30/30 | 33/33 | 39/40 | 24/24 |
|  | 2 | 40/40 | 34/34 | 34/35 | 33/33 | 35/35 |
| **RA (2 μM)** | 1 | 35/40 | 40/40 | 28/38 | 25/25 | 32/32 |
|  | 2 | 31/38 | 32/32 | 29/40 | 36/37 | 44/44 |
| **Ro (2 μM)** | 1 | 8/32 | 5/37 | 10/45 | 29/32 | 37/37 |
|  | 2 | 12/45 | 8/40 | 17/50 | 36/38 | 52/52 |

## Ethical declaration

All methods are reported in accordance with ARRIVE guidelines. All experiments and methods approved by the Animal Welfare and Ethical Review Board at the University of Exeter and University of Tokyo.

## Data Availability

The transcriptome data of *C. gigas* larvae treated with Ro41-5253 and atRA is desposited and publicly available from NCBI GEO accession GSE291194.

## Acknowledgements

This work was supported by the JSPS Overseas Long Term Fellowship to KS, the NC3Rs project grant (NC/X001121/1) to TK, and the JSPS KAKENHI grants (23244101 and 18H01323) to KE. We thank the Aquatic Resource Facility staff of the University of Exeter for maintenance and husbandry of animals. We also thank to Liz Williams and David Salamanca-Diaz for constructive comments on our manuscript and data, and to Paul O’Neill for bioinformatics support.

## References

Albalat, R. The retinoic acid machinery in invertebrates: ancestral elements and vertebrate innovations. Mol. Cell. Endocrinol. 313, 23–35 (2009).

Albalat, R. & Cañestro, C. Identification of Aldh1a and Cyp26 and RAR orthologs in protostomes pushes back the retinoic acid genetic machinery in evolutionary time to the bilaterian ancestor. Chem. Biol. Interact. 178, 188–196 (2009).

Anderson, D. T. Embryology and phylogeny in annelids and arthropods. Pergamon Press, Oxford (1973).

Aulehla, A. & Pourquié, O. Signaling gradients during paraxial mesoderm development. Harb. Perspect. Biol. 2, a000869 (2010).

Baratte, S., Andouche, A. & Bonnaud, L. Engrailed in cephalopods: a key gene related to the emergence of morphological novelties. Dev. Genes Evol. 217, 353–62 (2007).

Campo-Paysaa, F., Marlétaz F, Laudet, V. & Schubert, M. Retinoic acid signaling in development: tissue-specific functions and evolutionary origins. Genesis 46, 640–656 (2008).

Cather, J. N. Cellular interactions in the development of the shell gland of the gastropod, Ilyanassa. J. Exp. Zool. 166, 205–23 (1967).

Conway-Morris, S. The cuticular structure of the 495-Myr-old type species of the fossil worm Palaeoscolex, P. piscatorum. Zoological Journal of the Linnean Society 119, 69–82 (1997).

Créton, R., Zwaan, G. & Dohmen, R. Specific developmental defects in molluscs after treatment with retinoic acid during gastrulation. Dev. Growth Differ., 357–364 (1993).

Deguchi, R. Fertilization causes a single Ca2 þ increase that fully depends on Ca2 þ influx in oocytes of limpets (Phylum Mollusca, Class Gastropoda). Dev. Biol. 304, 652–663. (2007).

Dmetrichuk, J.M., Carlone, R.L. & Spencer, G. E. Retinoic acid induces neurite outgrowth and growth cone turning in invertebrate neurons. Dev. Biol. 294, 39–49 (2006).

Dzik, J. Early metazoan evolution and the meaning of its fossil record. In M. K. Hecht, R. J. MacIntyre, and M. T. Clegg (eds.), Evolutionary biology, pp. 339–386. New York (1993)

Germain P, Staels B, Dacquet C, Spedding M, Laudet V: Overview of nomenclature of nuclear receptors. Pharmacol Rev 2006, 58:685–704.

Grabherr, M. G. et al. Full-length transcriptome assembly from RNA-Seq data without a reference genome. Nat. Biotechnol. 29, 644–652 (2011).

Hashimoto, N., Kurita, Y. & Wada, H. Developmental role of dpp in the gastropod shell plate and co-option of the dpp signaling pathway in the evolution of the operculum. Dev. Bio. 366, 367–373 (2012).

Haas, B. J. et al. De novo transcript sequence reconstruction from RNA-seq using the Trinity platform for reference generation and analysis. Nat. Protoc. 8, 1494–1512 (2013). Deployment of regulatory genes during gastrulation and germ layer specification in a model spiralian mollusc Crepidula.

Perry KJ, Lyons DC, Truchado-Garcia M, Fischer AH, Helfrich LW, Johansson KB, Diamond JC, Grande C, Henry JQ. Dev Dyn. 244:1215-48. (2015)

Hinman VF, O’Brien EK, Richards GS, Degnan BM. Expression of anterior Hox genes during larval development of the gastropod Haliotis asinina. Evol Dev 5 508–21 (2003).

Holder, N., & Hill, I. Retinoic acid modifies development of the midbrain-hindbrain border and affects cranial ganglion formation in zebrafish embryos. Development 113, 1159–1170 (1991).

Holland, L. Z. Non-neural ectoderm is really neural: evolution of developmental patterning mechanisms in the non-neural ectoderm of chordates and the problem of sensory cell homologies. J. Exp. Zool. B Mol. Dev. Evol. 304, 304–323 (2005).

Holland, L. Z., Kene, M. & Holland, N. D. Sequence and embryonic expression of the amphioxus engrailed gene (AmphiEn): The metameric pattern of transcription resembles that of its segment-polarity homolog in Drosophila. Development 124, 1723–1732. (1997).

Iijima M, Takeuchi T, Sarashina I, Endo K. Expression patterns of engrailed and dpp in the gastropod Lymnaea stagnalis. Dev Genes Evol. 218, 237–51 (2008).

Jacobs, D. K. & Gates R. D. Developmental Genes and the Reconstruction of Metazoan Evolution – Implications of Evolutionary Loss, Limits on Inference of Ancestry and Type 2 Errors. Integrative & Comparative Biology v. 43 pp. 619–646 (2003).

Jacobs, D. K., Wray, C. G., Wedeen, C. J., Kostiken, R., Desalle, R., Staton, J. L., Gates, R. D. & Lindberg, D. R. Molluscan engrailed expression, serial organization, and shell evolution. Evol. Dev. 2, 340–347 (2000).

Kin, K., Kakoi, S. & Wada, H. Novel role for dpp in the shaping of bivalve shells revealed in a conserved molluscan developmental program. Dev. Biol. 329, 152–166 (2009).

Kniprath, E. Ontogeny of the molluscan shell field. Zoologica Scripta 10, 61–79 (1981).

Kudoh, T., Wilson, S. W. & Dawid, I. B. Distinct roles for Fgf, Wnt and retinoic acid in posteriorizing the neural ectoderm. Development 129, 4335–4346 (2002).

Lans, D., Wedeen, C. J. & Weisblat, D. A. Cell lineage analysis of the expression of an engrailed homolog in leech embryos. Development 117, 857–871 (1993).

Lowe, C. J., Wray, G. A. Radical alterations in the roles of homeobox genes during echinoderm evolution. Nature 16, 718–721. (1997)

Lambert DJ. Developmental Patterns in Spiralian Embryos Curr Biol. 20, R72–R77 (2010).

Malinda, K. M. & Ettensohn, C. A. Primary mesenchyme cell migration in the sea urchin embryo: distribution of directional cues. Dev. Biol. 164, 562–578 (1994).

Manzanares, M., et al. Conservation and elaboration of Hox gene regulation during evolution of the vertebrate head. Nature 408, 854– 857 (2000).

Marshall, H., et al. A conserved retinoic acid response element required for early expression of the homeobox gene Hoxb-1. Nature 370, 567–571 (1994).

Moshel, S. M., Levine, M. & Collier, J. R. Shell differentiation and engrailed expression in the Ilyanassa embryo. Dev. Genes Evol. 208, 135–141 (1998).

Nederbragt, A. J., van Loon, A. E. & Dictus, W. J. Expression of Patella vulgata orthologs of engrailed and dpp-BMP2/4 in adjacent domains during molluscan shell development suggests aconserved compartment boundary mechanism. Dev. Biol. 246, 341–55 (2002).

Nüsslein-Volhard, C. & E. Weischaus. Mutations affecting segment number and polarity in Drosophila. Nature 287, 795–801 (1980).

Ohta S, Noshita K, Kimoto K, Ishikawa A, Sato H, Shimizu K, Endo K. Possible roles of Wnt in the shell growth of the pond snail Lymnaea stagnalis. Sci Rep. 14:26488. (2024)

Patel, N. H., Kornberg, T. B. & Goodman, C. S. Expression of engrailed during segmentation in grasshopper and crayfish. Development 107, 201–212 (1989).

Perry KJ, Lyons DC, Truchado-Garcia M, Fischer AH, Helfrich LW, Johansson KB, Diamond JC, Grande C, Henry JQ. Deployment of regulatory genes during gastrulation and germ layer specification in a model spiralian mollusc Crepidula. Dev Dyn. 244, 1215–48. (2015).

Ramskold, L. & Hou, X. New early Cambrian animal and onychophoran affinities of enigmatic metazoans. Nature 351: 225–228 (1991).

Reijntjes, S., Blentic, A., Gale, E., & Maden, M. The control of morphogen signalling: regulation of the synthesis and catabolism of retinoic acid in the developing embryo. Dev. Biol. 285, 224–237 (2005).

Romero, R. & Bueno, D. Disto-proximal regional determination and intercalary regeneration in planarians, revealed by retinoic acid induced disruption of regeneration. Int. J. Dev. Biol. 45, 669–673 (2001).

Schilling, T. F., & Knight, R. D. Origins of anteroposterior patterning and Hox gene regulation during chordate evolution. Philos. Trans. R. Soc. Lond. B. Biol. Sci. 356 1599–1613 (2001).

Seaver, E. C. & Kaneshige, L. M. Expression of ‘segmentation’ genes during larval and juvenile development in the polychaetes Capitella sp. I and H. elegans. Dev. Biol. 289, 179–194 (2006).

Shimeld, S. M. Retinoic acid, hox genes and the anterior-posterior axis in chordates. BioEssays 18, 613–616 (1996).

Shimizu, K., Sarashina, I., Kagi, H. & Endo, K. Possible functions of dpp in gastropod shell formation and shell coiling. Dev. Genes Evol. 221, 59–68 (2011).

Shimizu, K., Iijima, M., Setiamarga, D. H. E., Sarashina, I., Kudoh, T., Asami, T., Gittenberger, E. & Endo, K. Left-right asymmetric expression of dpp in the mantle of gastropods correlates with asymmetric shell coiling. EvoDevo 4, 15. (2013).

Shimizu, K., Luo, Y-J., Satoh, N. & Endo, K. Possible co-option of engrailed during brachiopod and mollusc shell development. Biol. Lett. 13, 20170254. 10.1098/rsbl.2017.0254 (2017).

Vogeler, S., Galloway, T. S., Isupov, M. & Bean, T. P. Cloning retinoid and peroxisome proliferator-activated nuclear receptors of the Pacific oyster and in silico binding to environmental chemicals. PLoS ONE 12, e0176024. 10.1371/journal.pone.0176024 (2017).

Vachon, G., Cohen, B., Pfeifle, C., McGuffin, M. E., Botas, J. & Cohen, S. M. Homeotic genes of the bithorax complex repress limb development in the abdomen of the Drosphila embryo through the target gene *Distal-less*. Cell 71, 437–450 (1992).

Wada, H. Origin and evolution of the neural crest: a hypothetical reconstruction of its evolutionary history. Dev. Growth Differ. 43, 509– 520 (2001).

Wanninger, A. & Haszprunar, G. The expression of an engrailed protein during embryonic shell formation of the tusk-shell, *Antalis entalis* (Mollusca, Scaphopoda). Evol. Dev. 3, 312–21 (2001).

Wedeen, C. J., & Weisblat, D. A. Segmental expression of an engrailed-class gene during early development and neurogenesis in an annelid. Development 113, 805–814 (1991).

Wedeen, C. J., Kostriken, R. G., Leach, D., & Whitington, P. Segmentally iterated expression of an engrailed-class gene in the embryo of an Australian onychophoran. Dev. Genes Evol. 270, 282–286 (1997).

Yang, J., Martin, R. S., Lan, T., Hou, J. & Zhang, X. Articulated Wiwaxia from the Cambrian Stage 3 Xiaoshiba Lagerstätte. Scientific Reports 4 (2014).

Zamora, S., Sumrall, C. D. & Vizcaïno, D. Morphology and ontogeny of the Cambrian edrioasteroid echinoderm Cambraster cannati from western Gondwana. Acta Palaeontol. Polonica (2013).

Zhao, F., Bottjer, D. J., Hu, S., Yin, Z. & Zhu, M. Complexity and diversity of eyes in early Cambrian ecosystems. Sci. Rep. 3, 2751 (2013).

